# TEITbase: a database for transposable element (TE)-initiated transcripts in human cancers

**DOI:** 10.1101/2025.05.19.654796

**Authors:** Yun Zhang, Jianqi She, Xueyan Hu, Yueqi Jin, Changyu Tao, Minghao Du, Ence Yang

## Abstract

Transposable elements (TEs) are abundant and play a crucial regulatory role in the human genome. Functioning as alternative promoters, TEs can be reactivated to produce TE-initiated transcripts. In cancer, TE-initiated transcription may upregulate oncogene expression and generate novel tumor-specific antigens, which could serve as potential targets for immunotherapy. However, there remains a lack of comprehensive databases that systematically investigate TE-initiated transcription in cancer. To address this gap, we developed a deep learning-based method and identified 38,995 TE-initiated transcripts across 33 cancer types from The Cancer Genome Atlas. Among these, 6,203 were tumor-specific expressed and strongly associated with the tumorigenesis. Using these annotations, we created TEITbase (http://teitbase.medbioinfo.org/), a user-friendly database that provides researchers with tools to conduct various downstream analyses and investigations for TE-initiated transcripts, including 546 novel onco-exaptation events. The establishment of TEITbase provides new insights into transcriptional reprogramming in cancers and enables further investigation into the potential roles of TE-initiated transcripts in cancer diagnostics and therapeutic strategies.

## Introduction

Transposable elements (TEs) are mobile genetic elements that are widespread in eukaryotic genomes (1). Approximately 45% of the human genome is derived from various classes of TEs, such as LINEs (long interspersed nuclear elements), SINEs (short interspersed nuclear elements), LTRs (long terminal repeats), and DNA transposons (2,3). In humans, only a small proportion of TEs, including active human-specific L1 (L1HS) elements, remain capable of transposition and inducing genetic changes (4,5). The majority of TEs in the human genome are incomplete and no longer transposable (6). However, TE-derived sequences retain significant regulatory potential, acting as promoters, enhancers, splicing sites, and more, playing crucial roles in both physiological and pathological processes (6–9).

As potential threats to human genome, most TEs have been kept inactive in somatic cells by mutational events and epigenetic repression (6,10). However, accumulating evidence suggests that certain TEs are transcriptionally reactivated in human cancers as a result of epigenetic dysregulation (11,12). Acting as alternative promoters, TEs can enhance their own transcription or regulate the expression of neighboring genes, leading to the production of TE-initiated transcripts with TE-derived transcription start sites (TSSs) (13). In cancer, TE-initiated transcription can upregulate the expression of oncogenes, a phenomenon known as onco-exaptation (14–16). Notable examples include Alujb-*LIN28B*, L1-*FGGY* and HERVH-*CALB1* (15,17,18). Furthermore, TE-initiated transcription may produce novel protein isoforms and tumor-specific antigens, which could serve as potential targets for immunotherapy (19,20). For instance, the tumor-specific membrane protein GABRG2, transcribed from the L1PA2 promoter, can generate aberrant epitopes on the extracellular surface of cancer cells (12). Moreover, endogenous retrovirus (ERV) envelope glycoproteins have been identified as a dominant anti-tumor antibody target and ERV-reactive antibodies exhibit anti-tumor activity that extends survival in mouse models (21).

Given the recognized role of TEs in tumor progression, several databases, including ERVcancer and CancerHERVdb, have been developed to investigate TE expression in human cancers (22,23). However, these databases only provide TE expression data at the TE family level, losing locus information and failing to distinguish between TE-initiated transcription and TE exonization. Recently, Gu et al. established TE-TSS, a database compiling 2,681 RNA sequencing datasets that identifies 5,768 human TE-derived TSSs and 2,797 mouse TE-derived TSSs (24). Nevertheless, due to the repetitive nature of TEs and the limitations of short-read RNA-seq, TE-TSS cannot provide accurate annotations of TE-initiated transcripts. Deep learning techniques, which can automatically extract features from DNA sequence information, have been widely applied to predict TSSs (25,26). By integrating RNA-seq data with DNA sequence information, convolutional neural networks could accurately predict TSSs for each assembled transcript, thereby enabling the identification of TE-initiated transcripts. However, there remains a lack of databases that specifically annotate TE-initiated transcripts in human cancers.

In this work, we developed the TEITbase (http://teitbase.medbioinfo.org/), an integrated database that systematically studies the TE-initiated transcription across 33 cancer types from The Cancer Genome Atlas (TCGA), offering data at both the TE family and transcript levels. To identify TE-initiated transcripts, we first developed a deep learning-based method to predict transcription start sites (TSSs) for each assembled transcript, utilizing both DNA sequences and short-read RNA sequencing data. Subsequently, we applied this method to 10,079 tumor samples from TCGA, and identified 38,995 TE-initiated transcripts. These annotations enable TEITbase to provide researchers with the tools to conduct various downstream analyses, such as promoter activity of TE families, tumor-specific TE-initiated transcripts, differential expression analysis, stage analysis, and survival analysis. We believe that TEITbase will serve as a valuable resource for further understanding the role of TE-initiated transcription in epigenetically dysregulated tumor tissues and will be instrumental in investigating novel tumor-specific antigens that could potentially be therapeutically targeted.

## Results

### A deep learning-based method for the identification of TE-initiated transcript

Due to the repetitive nature of TEs and the limitations of short-read RNA-seq, identifying TE-derived TSSs based on RNA-seq coverage is not sufficiently accurate. To overcome the challenge, we developed a deep learning-based method that integrates both DNA sequence and RNA-seq coverage information. Using one-hot encoding and convolutional neural network, we predicted TSSs for each assembled transcript and overlapped them with TE annotations to identify TE-initiated transcripts (Fig. 1A).

**Fig. 1.**
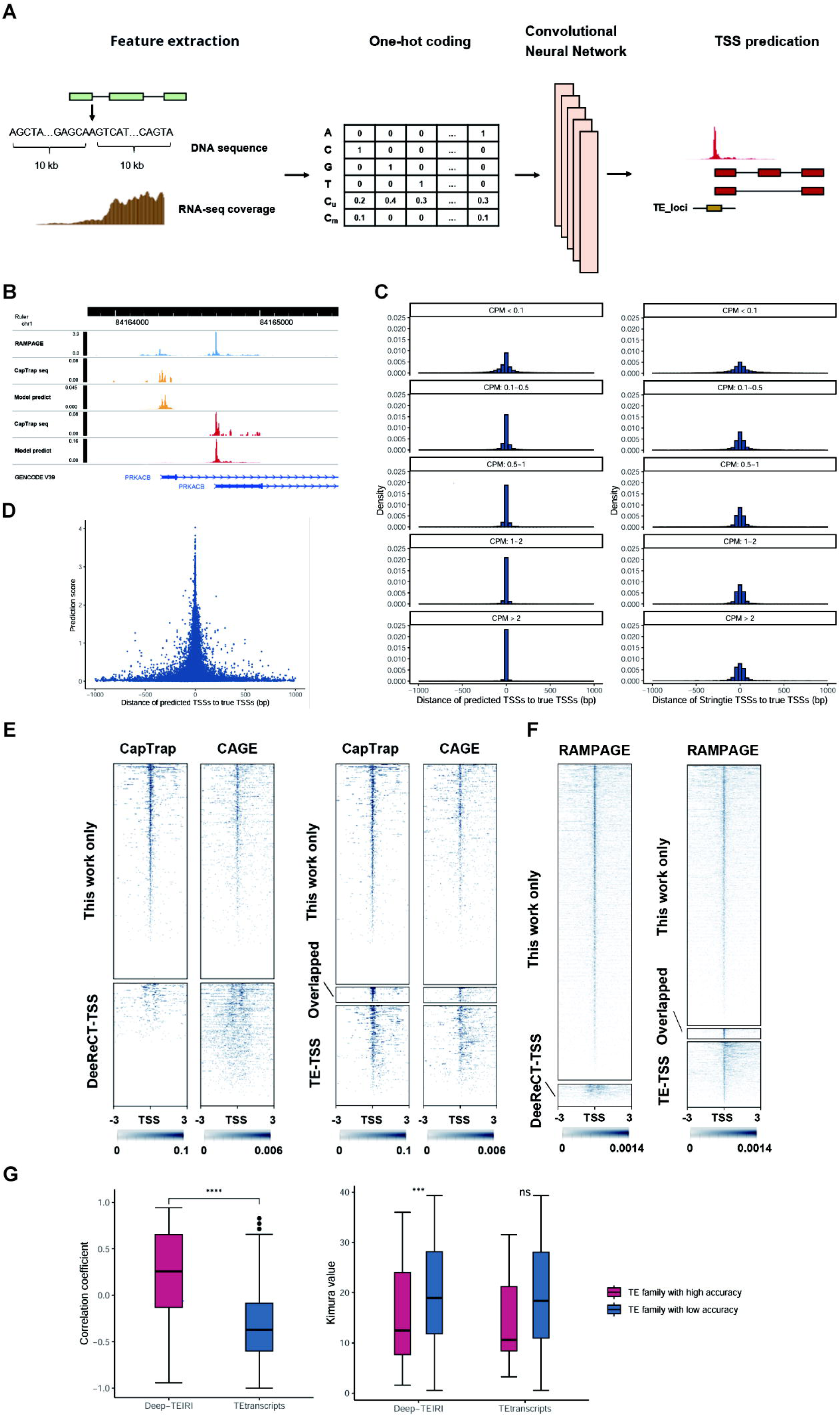
Detection of TE-initiated transcripts. (**A**) Pipeline designed to identify TE-initiated transcripts by using a deep learning model. (**B**) RAMPAGE and CapTrap-seq signals of the *PRKACB* gene, along with TSS predictions of our method. (**C**) The distances to the true TSSs of predicted TSSs from our method (left) and Stringtie (right). (**D**) Prediction score and distance of predicted TSSs to the true TSSs. (**E**) CapTrap-seq and CAGE signals around the TE-derived TSSs identified by our method or DeeReCT-TSS (left) and those identified in this work, TE-TSS database, or both (right) in H1 cell lines. (**F**) RAMPAGE signals around the TE-derived TSSs identified by our method or DeeReCT-TSS (left) and those identified in this work, TE-TSS database, or both (right) in testis tissues. (**G**) Spearman correlation coefficient between CAGE data and the quantitative results from our method or TEtranscripts (left). The Kimura value of high-accuracy TE families and others (right).

To train our model, we downloaded CapTrap-seq datasets along with matched short-read RNA-seq datasets (27). Unlike CAGE and RAMPAGE, CapTrap-seq combines the cap-trapping strategy with long-read sequencing technologies, enabling the identification of both TSSs and full-length transcripts. Using these CapTrap-seq datasets for training, our model can predict TSSs at the transcript level. For example, *PRKACB* has alternative promoters in brain tissue. While RAMPAGE identifies only two TSS signals, it cannot assign them to specific transcripts. In contrast, both CapTrap-seq and our model can identify the TSS for each transcript (Fig. 1B).

Compared to StringTie, which relies solely on RNA-seq coverage to determine TSS, our model significantly enhances the accuracy of TSS prediction (*P* = 4.6 × 10^−4^, Fig. 1C, Table S1). In the validation set, our model accurately identified an average of approximately 26.7% of TSSs, with about 84.7% of TSSs exhibiting identification errors within 100 bp, a notable improvement over StringTie’s performance of 2.4% and 70.6%, respectively. As expected, our model performed significantly better in predicting highly expressed transcripts (Fig. 1C, *P* = 1.2 × 10^−5^). For transcripts with counts per million (CPM) greater than 2, an average of approximately 71.0% of TSSs were accurately identified by our model, with around 99.2% of TSSs having identification errors within 100 bp. This is substantially better than for low-expressed transcripts (CPM < 0.1), which had 9.1% and 72.7% accuracy within 100 bp, respectively. Furthermore, the prediction score of our model was significantly and negatively correlated with TSS prediction errors (Fig. 1D, Spearman’s *r* = –0.50, *P* < 2.2 × 10^−16^), enabling us to improve TSS identification accuracy by setting an appropriate prediction score threshold.

Through the application of the deep learning model and strict criteria, our method significantly outperforms DeeReCT-TSS in identifying TE-TSS (Table S2, *P* = 0.029) (26). In the H1 cell line, our method identified 954 TE-TSS, of which 626 (65.6%) were supported by CapTrap-seq data. In contrast, DeeReCT-TSS identified only 390 TE-TSS, with just 12 (3.1%) supported by CapTrap-seq data (Fig. 1E). When compared to the TE-TSS database (24), our method identified 64 previously annotated TE-TSS and 908 novel TE-TSS, of which 580 (63.9%) were supported by CapTrap-seq data (Fig. 1E). To exclude the potential impact of model overfitting on TE-TSS identification, we downloaded short-read RNA-seq data from 18 testis tissues as an external validation dataset (28), with RAMPAGE data serving as the gold standard. Our method identified 3,979 TE-TSS, of which 2,909 (73.1%) were supported by RAMPAGE data, while DeeReCT-TSS identified only 286 TE-TSS, with just 107 (37.4%) supported by RAMPAGE data (Fig. 1F). Compared to the TE-TSS database, our method identified 157 previously annotated TE-TSS and 3,822 novel TE-TSS, of which 2,752 (72.0%) were supported by RAMPAGE data (Fig. 1F). These results suggest that we have developed a deep learning-based method that can accurately identify TE-initiated transcripts using multiple biological replicates of short-read RNA-seq data, significantly outperforming existing methods.

Due to the relatively accurate annotation of TE-initiated transcripts, our method effectively distinguishes between TE-initiated transcription and TE exonization, enabling the measurement of promoter activity in TE families. To evaluate the quantitative performance at the family level, we downloaded short-read RNA-seq data and corresponding CAGE data (used as the gold standard for quantitative evaluation) from six cancer cell lines (A549, Hela, HepG2, K562, MCF-7, and SK-N-SH) from the ENCODE database (29). Our method demonstrated significantly better consistency with CAGE data compared to existing TE family expression quantification methods, TEtranscripts (Fig. 1G, *P* < 2.2 × 10^−16^). Among all TE families, the median Spearman correlation coefficient between the quantitative results from our method and CAGE data across these six cell lines was 0.26, while TEtranscripts had a value of –0.37. Furthermore, our method identified 105 TE families with high accuracy (Spearman’s *r* ≥ 0.7), whereas TEtranscripts identified only 6. Notably, the age of these 105 high-accuracy TE families (measured by the evolutionary distance to ancestral TEs: Kimura value) was significantly lower than that of other TE families (Fig. 1G, *P* = 9.2 × 10^−4^), which may be related to the higher retention of promoter activity in younger TEs.

### Genome-wide identification of TE-initiated transcripts in pan-cancers

In 10,079 tumor samples across 33 cancer types from the TCGA, we identified 38,995 TE-initiated transcripts (Table S3), of which 34,670 were not annotated in GENCODE.v46 (30). The expression of these TE-initiated transcripts was associated with the activation of 39,703 TE loci, including 11,857 SINE elements, 11,758 LTR elements, 10,911 LINE elements, and 2,580 DNA transposons (Fig. 2A). These active TEs contributed to the expression of 14,453 neighboring genes, including 9,315 protein-coding genes and 4,771 lncRNA genes (Fig. 2B), suggesting that TEs play a widespread role in regulating gene expression in tumor tissues.

**Fig. 2.**
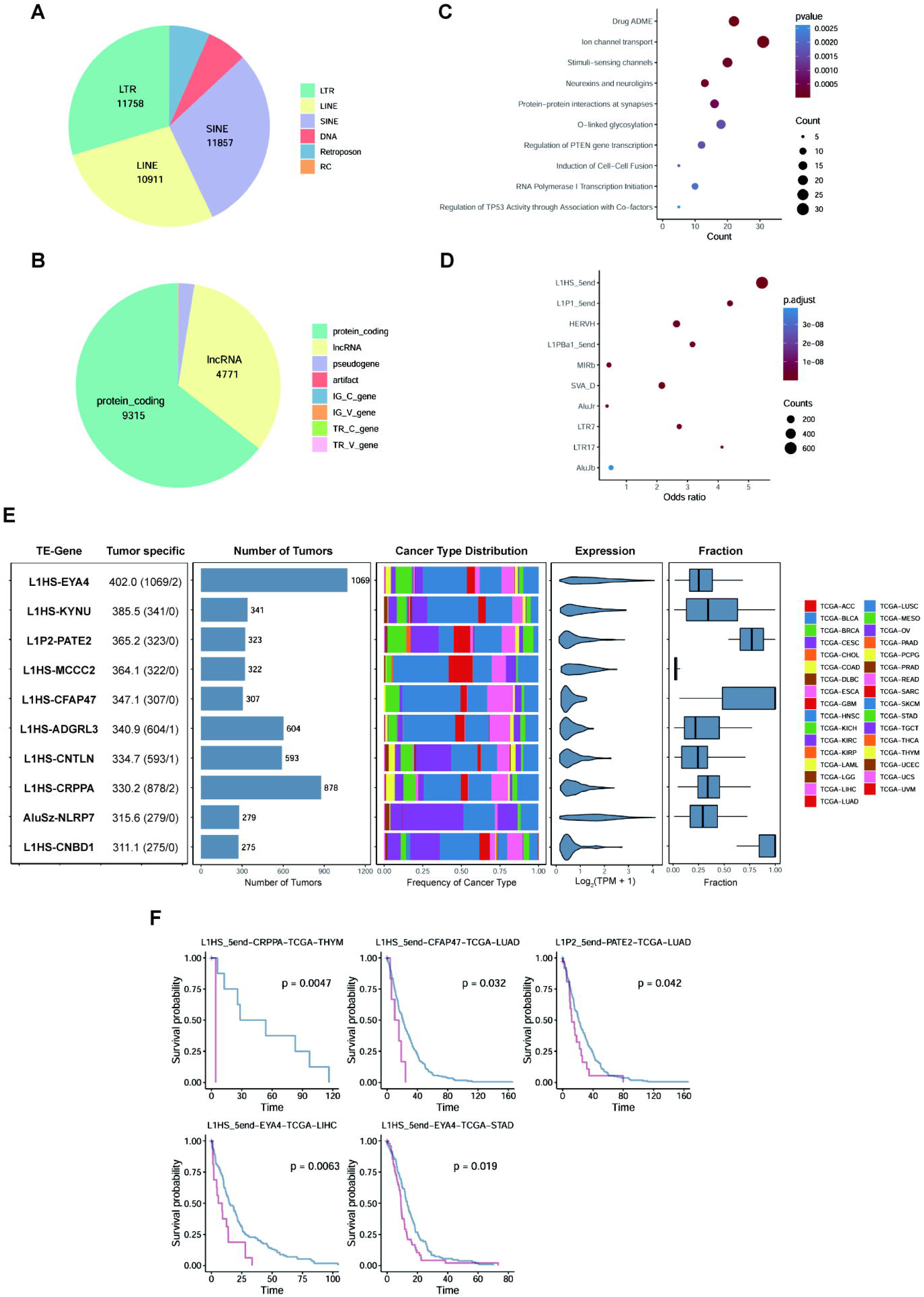
Tumor-specific TE-initiated transcription promotes tumorigenesis. (**A**) The distribution of the activated TE loci across the TE classes. (**B**) The distribution of the neighboring genes across the gene type. (**C**) The reactome pathway enrichment analysis of neighboring genes associated with the tumor-specific TE-initiated transcripts. (**D**) The TE family enrichment analysis of tumor-specific TE-initiated transcripts. (**E**) The 10 TE-initiated transcripts with the highest tumor-specific scores are presented. The left-most panel shows the TE-gene along with the tumor-specific score. The next two panels display the number of tumor samples in which each TE-initiated transcript is expressed and the distribution across cancer types. The final two panels show the expression levels of the TE-initiated transcript and the percentage of total oncogene expression contributed by TE-initiated transcription. (**F**) Kaplan–Meier plots demonstrating overall survival associated with the expression of TE-initiated transcripts.

Tumor-specific expressed transcripts play a critical role in tumorigenesis and represent important sources of tumor-specific antigens, which may provide new targets for cancer immunotherapy. To investigate this, we performed the expression analysis of TE-initiated transcripts in 10,079 tumor samples, 740 tumor-adjacent samples and 10,620 non-testicular normal tissue samples. Tumor-specific scores were calculated, identifying 6,203 tumor-specific TE-initiated transcripts (scores ≥ 8, Table S4). These activated tumor-specific TE-initiated transcripts were associated with the expression of 1,532 adjacent protein-coding genes. As anticipated, enrichment analysis revealed significant overrepresentation in tumorigenesis-related pathways, including regulation of *PTEN* gene transcription and TP53 activity (Fig. 2C, Table S5). Among the 702 oncogenes annotated in the ONGene database (31), we found that TEs upregulated the expression of 333 oncogenes (a process known as onco-exaptation), generating 696 TE-initiated transcripts (Table S6), including previously reported onco-exaptations such as AluJb-*LIN28B*, L1HS-*SYT1*, L1HS-*FGGY*, and L1HS-*MET* (15,18,32). Compared to a recent similar study (15), we identified 150 reported and 546 novel onco-exaptation events, with 263 (48.2%) supported by CAGE data from at least one cancer cell line. Among them, AluJr-*AKT2* was expressed across all cancer types and detected in 619 tumor samples, further highlighting the universality of onco-exaptation events. Furthermore, TE family enrichment analysis revealed significant overrepresentation of young families, including L1HS, L1P1, and HERVH (Fig. 2D, Table S7). These evolutionarily young TE families retain regulatory sequences that are typically suppressed in somatic cells but are reactivated in the epigenetically dysregulated tumor tissues, driving the TE-initiated transcription and potentially promoting tumor progression.

To illustrate, we analyzed the 10 TE-initiated transcripts with the highest tumor-specific scores and found that 8 were initiated by the L1HS family (Fig. 2E). These 10 TE-initiated transcripts were expressed in 275 to 1,069 tumor samples but only in 1 or 2 normal tissue samples, indicating their high tumor specificity. Moreover, these activated TEs were critical in regulating the expression of adjacent genes, with L1HS-*CNBD1*, L1HS-*CFAP47*, and L1P2-*PATE2* serving as the primary transcripts for their respective genes. Furthermore, *EYA4* and *NLRP7* have been shown to promote cell proliferation, invasion, and migration in various cancers (33,34), suggesting that the expression of L1HS-*EYA4* and AluSz-*NLRP7* may have significant oncogenic effects. Survival analysis further indicated that the expression of L1HS-*EYA4*, L1HS-*CRPPA*, L1HS-*CFAP47*, and L1P2-*PATE2* was associated with poor prognosis in at least one cancer type (Fig. 2F). In conclusion, these highly tumor-specific TE-initiated transcripts may play a role in tumorigenesis and hold potential as biomarkers for tumor diagnosis and prognosis.

### Content and usage of TEITbase

Based on the identification of TE-initiated transcripts across various cancer types, we have developed a free, user-friendly website, TEITbase (http://teitbase.medbioinfo.org/), which provides expression data for 1,027 TE families and 38,995 TE-initiated transcripts (Fig. 3A). The website consists of several main pages: Home, Browse, Analyze, Statistics, Download, Help, and Contact. TEITbase offers an intuitive interface and a range of core functionalities, enabling researchers to explore and understand the role of TE-initiated transcription in cancers.

**Fig. 3.**
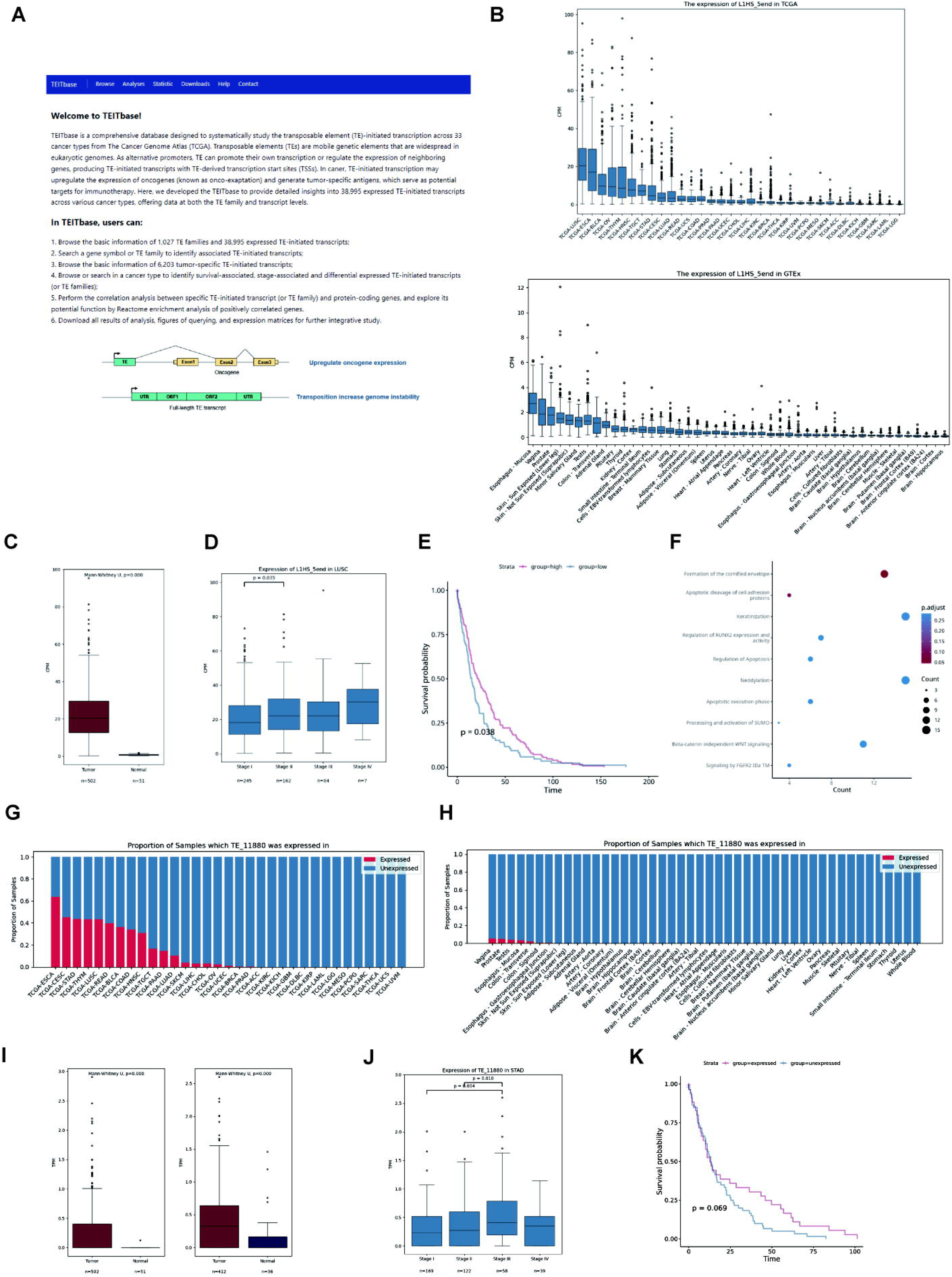
Applications of TEITbase. (**A**) The homepage of TEITbase. (**B**) Box plot showing the promoter activity of L1HS in TCGA tumors (top) and GTEx normal tissues (bottom). (**C**) Box plot showing the promoter activity of L1HS in LUSC and adjacent normal tissues. (**D**) Box plot illustrating the promoter activity of L1HS across different stages of LUSC. (**E**) Kaplan–Meier plots showing overall survival associated with the promoter activity of L1HS in LUSC. (**F**) The reactome pathway enrichment analysis of genes positively correlated with the promoter activity of L1HS in LUSC. (**G**, **H**) Proportion of TCGA tumor samples (**G**) and GTEx normal tissue samples (**H**) expressing TE_11880. (**I**) Box plot showing the expression level of TE_11880 in LUSC, STAD and adjacent normal tissues. (**J**) Box plot illustrating the expression level of TE_11880 across different stages of STAD. (**K**) Kaplan–Meier plots showing overall survival associated with the expression of TE_11880 in COAD.

The Browse page is divided into two sections: TE Family and TE-initiated Transcripts. Users can query TE family entries by TE family, TE superfamily, and species. The query results display the TE family, Dfam database ID, TE superfamily, TE classification, species, consensus sequence length, Kimura value, and tissue-specificity score (Tau index). Users can also query TE-initiated transcripts using TEITbase database ID, TE family, and gene. The results for these queries include the TEITbase database ID, TE family, TE superfamily, TE classification, chimeric gene, transcript classification, coding types, tumor specificity score, and TSS location. According to the GENCODE v46 annotation, TE-initiated transcripts are classified into three categories: annotated, chimeric (with a novel splice site at the 5’ end), and intergenic (without known splice sites). Based on predicted open reading frames (ORFs), TE-initiated transcripts are further categorized into six types: annotated, 5’ truncated, 5’ chimeric, chimeric truncated, other coding types, and non-coding. The Analyses page allows users to query tumor-specific TE-initiated transcripts, as well as survival-associated, stage-associated, and differentially expressed TE-initiated transcripts (or TE families). Additionally, the Statistics page provides the basic information of TEITbase, such as the number of TE-initiated transcripts. The Help page includes a user guide for the database, while the Download page offers links to download all annotation information, expression matrices, and analysis results.

Using the functions of TEITbase, we investigated the dysregulation of the L1HS family in cancer. We found that lung squamous carcinoma (LUSC), esophageal carcinoma (ESCA), and bladder urothelial carcinoma (BLCA) exhibit higher L1HS promoter activity, while normal tissues show lower activity (Fig. 3B). Differential expression analysis revealed significant activation of the L1HS promoter in 16 out of 21 cancer types, with the most pronounced increase in LUSC (Fig. 3C, *P* < 2.2 × 10^−16^). In LUSC, L1HS promoter activity in stage II patients was significantly higher than in stage I patients (Fig. 3D, *P* = 0.038). Survival analysis indicated that in 5 out of 33 cancer types, including liver hepatocellular carcinoma (LIHC), abnormal activation of the L1HS promoter was associated with poorer tumor prognosis. Conversely, activation of the L1HS promoter is significantly correlated with better tumor prognosis in LUSC (Fig. 3E), adrenocortical carcinoma (ACC), and mesothelioma (MESO). To explore how L1HS promoter activation improves prognosis, we conducted a correlation analysis between L1HS promoter activity and gene expression, followed by enrichment analysis of genes positively correlated with its expression. We discovered that in LUSC, activation of the L1HS promoter was linked to apoptosis-regulating signaling pathways (Fig. 3F), suggesting that active transposition of L1HS may disrupt tumor genome stability, leading to cancer cell apoptosis (35).

By utilizing the ‘Tumor-specific transcripts’ function of TEITbase, we investigated the role of TE_11880 (L1HS-*IRF1*) in cancers. TE_11880 is initiated by L1HS with a tumor-specific score of 32.5 and is chimeric with *IRF1*, a transcriptional regulator and tumor suppressor that activates genes involved in antitumor immunity (36). These findings suggest that the activation of TE promoters in cancer tissues may also confer benefits to the host. Further details about TE_11880, including its sequence and other relevant information, can be accessed in the TEITbase database. TE_11880 is silent in most normal tissue samples but activated in several cancer types (Fig. 3G, H). We selected specific cancer types to explore the differential expression, stage, and survival associations of TE_11880. Our differential expression analysis revealed that TE_11880 expression was primarily restricted to LUSC and stomach adenocarcinoma (STAD), with no expression in adjacent normal tissues (Fig. 3I). In STAD, TE_11880 expression in stage III patients was significantly higher than in both stage I and stage II patients (Fig. 3J). Additionally, TE_11880 expression is associated with a favorable prognosis in colon adenocarcinoma (COAD), despite not reaching statistical significance (*P* = 0.069) (Fig. 3K), suggesting its potential prognostic value.

## Discussion

The activation of TE promoters can significantly impact the transcriptomic and proteomic profiles of cancer (20). Recent studies suggest that TE-initiated transcription may be linked to tumorigenesis and could generate tumor-specific antigens, offering potential targets for immunotherapies (12,15). However, the widespread use of short-read RNA sequencing technologies poses challenges due to the loss of TSS information. To address this, we developed a deep learning-based approach that predicts TSSs and accurately identifies TE-initiated transcripts, enabling the creation of a specialized database, TEITbase. TEITbase offers abundant data resources and various analytical tools, allowing users to explore the roles of TE-initiated transcripts in cancer progression. Importantly, our findings have identified over 6000 tumor-specific TE-initiated transcripts, with the L1HS family being notably enriched. As the youngest L1 family in humans, the L1HS family retains transposable ability and may play a significant role in tumorigenesis (35).

Due to its powerful model fitting capabilities, deep learning has become widely applied to predicting key features from DNA sequences such as gene expression, RNA editing, TSS, alternative splicing, and polyadenylation (25,37–40). Several CNN-based tools have been developed for the accurate prediction of promoters and TSS, including Puffin, TSSPlant, TransPrise, and DeeReCT-TSS (25,26,41,42). Among them, DeeReCT-TSS accurately identifies active TSS across the genome by integrating DNA sequences and RNA-seq coverage information. However, DeeReCT-TSS uses CAGE-seq data to train the model, which loses relevant transcript structure information and cannot assign identified active TSS to specific transcripts. Besides, reads derived from TE sequences often produce ambiguous alignments, leading to errors in RNA-seq coverage estimation and reducing the accuracy of the identification of TE-derived TSS (43). To address this, we downloaded CapTrap-seq data to train the model (27), enabling the prediction of TSS for each transcript, and applied strict criteria to identify TE-initiated transcript. Our method significantly outperforms DeeReCT-TSS in identifying TE-derived TSS, providing a new tool to investigate TE-initiated transcription.

In summary, TEITbase is a user-friendly database that provides valuable insights and tools for investigating TE-initiated transcription in tumorigenesis, as well as exploring the potential roles of these transcripts in cancer diagnostics and therapeutic strategies. While it already offers a comprehensive data collection, there are still opportunities for optimization and further development. Future versions will incorporate TE exonization, which results from noncanonical splice junctions between exons and TEs, serving as a source of tumor-specific antigens (8). With ongoing advancements in full-length long-read RNA-seq technologies (16), TEITbase will integrate publicly available long-read RNA-seq data to capture the full length of TE-initiated and TE-exonization transcripts. These updates are expected to further enhance TEITbase’s utility and broaden its applicability across various research fields.

## Methods

### Data collection

We downloaded short-read RNA-seq data from 10,819 samples across 33 cancer types from the The Cancer Genome Atlas (TCGA, phs000178.v11.p8) (44). Additionally, we obtained RNA-seq data from 10,620 samples across 45 human body sites from the GTEx project (by dbGaP, study accession: phs000424.v8.p2) (45). Short-read RNA-seq and CapTrap-seq datasets were downloaded from the ENCODE project and the ArrayExpress repository (E-MTAB-13063) (27,29). Ten samples of these samples were used as the training set (H1 cell line and heart), and three samples were used as the validation set (endodermal cells and brain). Short-read RNA-seq and RAMPAGE datasets of testis were download from the Gene Expression Omnibus database (GSE135791) and the ENCODE project (28). Short-read RNA-seq and CAGE datasets of six cancer cell lines were download from the ENCODE project. GENCODE Version 46 was download from https://www.gencodegenes.org/human/ and used as the transcript reference (30). TE annotations and consensus sequences of TE family were downloaded from the Dfam database (https://www.dfam.org/releases/Dfam_3.8/) (46).

### Deep learning model for TSS prediction

CapTrap-seq and matched short-read RNA-seq datasets were used to train the model. For each 3’ end of the first exon, the 10 kb DNA sequence on either side, along with the corresponding RNA-seq coverage (including both unique-mapping and multi-mapping reads), was extracted and used as input to the model. The corresponding TSS signal from the CapTrap-seq data served as the model’s label. Our model is based on the architecture of the Puffin-D model, a convolutional neural network specifically designed for TSS prediction using DNA sequences (25). To enable TSS prediction for each sample, we adjusted the model to extract information from both the DNA sequence and RNA-seq coverage. To address the imbalance between positive and negative samples during model training, we employed the KL divergence loss function:

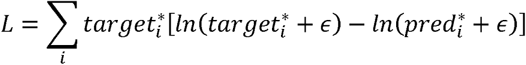

The *target* represents the true TSS signal at position *i*, *pred* represents the TSS signal predicted by the model, and □ is a constant set to *e*^−10^.

The model utilized the Adam optimizer for parameter optimization, with a learning rate of 0.005, a batch size of 64, and a maximum of 10 training epochs. Model construction and training were carried out using PyTorch. The distance between the predicted and actual TSS was used to evaluate the model’s performance. The example of an alternative promoter was visualized on WashU Epigenome Browser (47).

### Pipeline for the detection of TE initiated transcripts

First, RNA-seq data were aligned to the reference human genome (GRCh38) using STAR v2.7.10b (48). Sambamba was employed to remove PCR duplicates, and deepTools was used to extract RNA-seq coverage (49,50). StringTie v2.1.7 was used to assemble the reads into full-length transcripts (stringtie –m 200 –c 1 –t) (51). The TSS for each assembled transcript was then predicted using the trained deep learning model. These assembled transcripts across different samples were subsequently merged with a custom script. Briefly, the first exons of multi-exon transcripts were extracted. Following this, first exons overlapping with other internal exons were excluded, as they may have resulted from RNA degradation. Then, for each first exon, the TSS was corrected according to the score calculated as follows (52):

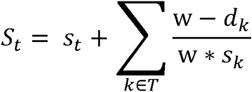

The *S* is the weight score for position *t*; *s* represents the number of samples supporting the position as a TSS of the first exon; *w* is the window size used to calculate the weight score; *T* is the set of all positions within a distance from *t* that is less than the window size; and *d* is the distance between positions *t* and *k*.

Corrected TSSs overlapping with TEs were retained, and the full-length transcripts of TE-initiated transcripts were merged using a reference-guided approach across samples. The count of junction reads supporting the first exon and the expression of TE-initiated transcripts at the transcript level were then calculated using StringTie. TE-initiated transcripts with at least one junction read supporting the first exon were considered expressed in the sample. The count of junction reads supporting the first exon of TE-initiated transcripts from the same TE family was summed, and the total read count mapped to the GRCh38 genome was used for normalization to obtain TE promoter activity at the family level.

### Computational validation of TE-derived TSSs and TE promoter activity

To evaluate the precision of TE-derived TSSs identified by our pipeline, we used CapTrap-seq or RAMPAGE data as the gold standard. A TE-derived TSS with supporting reads within 100 bp was considered positive. Plots of the CapTrap-seq and RAMPAGE signals surrounding the TSSs were generated using deepTools. To assess the precision of TE promoter activity calculated by our pipeline, the Spearman correlation with CAGE data was calculated.

### Annotation and expression analysis of TE-initiated transcripts

Identified TE-initiated transcripts were annotated with Dfam database by using BEDTools v2.31.1 (46,53). Based on the relationship with reference transcripts, TE-initiated transcripts were classified into three categories: annotated, chimeric, and intergenic. Coding potential of TE-initiated transcripts was calculated by CPC2 (54). For coding TE-initiated transcripts, we subsequently compared predicted CDS with annotated CDS using BLAST 2.14.1 (55). Then the coding TE-initiated transcripts were annotated into five classes: annotated, 5’ chimeric normal, 5’ chimeric truncated, 5’ truncated and novel. Tumor-specific scores for each TE-initiated transcript were calculated as follows:

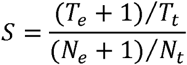

The *S* represents the tumor-specific score for the TE-initiated transcript. *T_e_* represents the number of tumor samples expressing the TE-initiated transcript, *N_e_* represents the number of tumor-adjacent or normal tissue samples expressing the TE-initiated transcript (excluding testicular tissue), and *T_t_* and *N_t_* represent the total number of tumor and normal samples, respectively. A TE-initiated transcript with a tumor-specific score greater than eight was considered tumor-specific expressed.

The reactome pathway enrichment analysis of gene-chimeric TE-initiated transcripts was conducted by the ReactomePA package (56). The TE family enrichment analysis was conducted using a two-tailed Fisher’s exact test. All *P*-values were corrected using the Benjamini-Hochberg (BH) procedure.

### Clinically associated TE families or TE-initiated transcripts

The Wilcoxon rank-sum test was used to assess statistical differences in the expression levels of TE families and TE-initiated transcripts between tumor and adjacent normal tissues. The Kruskal-Wallis test was applied to evaluate statistical differences in the expression levels of TE families and TE-initiated transcripts across patients with different tumor stages. The survdiff function from the survival package was used to perform the Log-Rank test, comparing differences between two survival curves, and calculating the hazard ratio (HR) and *P*-value. All *P*-values were adjusted using the Benjamini-Hochberg (BH) procedure.

### Construction of the TEITbase website

The backend application and frontend interface of the TEITbase website were developed using the Django framework, with Gunicorn serving as the WSGI HTTP server to execute the Django code. NGINX was employed as a reverse proxy server to forward client requests to Gunicorn. To ensure system stability, Supervisor was used to monitor the operation of the Gunicorn and Django processes. The frontend interface was designed and implemented using HTML, JavaScript, and CSS. Additionally, annotations for TE families and TE initiated transcripts were securely stored in an SQLite3 database.

## Declarations

### Ethics approval and consent to participate

Not applicable.

### Availability of data and materials

Details about data analyzed in this study were included in the Methods section. The modified pipeline, together with the deep learning model for TSS prediction developed in this study are available in (https://github.com/yunzhang1998/Deep-TEIRI).

## Author contributions

E.Y. conceived and designed the study. Y.Z., J.Q.S., X.Y.H., Y.Q.J., C.Y.T., and M.H.D. analyzed the data. Y.Z., J.Q.S., M.H.D., and E.Y. interpreted the data and wrote the manuscript. All authors approved the final version of the manuscript.

## Acknowledgments

We would like to express our gratitude to the TCGA and GTEx committees for their valuable contributions. We also acknowledge the ENCODE Consortium and the ENCODE production laboratories.

## Additional files

**Additional file 1: Table S1-7.** Supplementary tables (Table S1-S7, XLSX).

